# *ParS*-independent recruitment of the bacterial chromosome-partitioning protein ParB

**DOI:** 10.1101/2021.11.02.466941

**Authors:** Miloš Tišma, Maria Panoukidou, Hammam Antar, Young-Min Soh, Roman Barth, Biswajit Pradhan, Jaco van der Torre, Davide Michieletto, Stephan Gruber, Cees Dekker

## Abstract

The ParAB*S* system plays an essential role in prokaryotic chromosome segregation. After loading at the *parS* site on the genome, ParB proteins rapidly redistribute to distances of ~15 kb away from the loading site. It has remained puzzling how this large-distance spreading can occur along DNA that is loaded with hundreds of proteins. Using single-molecule *in vitro* visualization, we here show that, unexpectedly, ParB can load onto DNA independently and distantly of *parS,* whereby loaded ParB molecules are themselves able to recruit additional ParB proteins from bulk. Strikingly, this recruitment can occur *in-cis* but also *in-trans* whereby, at low tensions within the DNA, newly recruited ParB can bypass roadblocks as it gets loaded to spatially proximal but genomically distant DNA regions. The data are supported by Molecular Dynamics simulations which also show that cooperative ParB-ParB recruitment enhances spreading. *ParS-*independent recruitment explains how ParB can cover substantial genomic distance during chromosome segregation which is vital for the bacterial cell cycle.

---

Accurate chromosome segregation is crucial for a stable transmission of genetic material through each cell cycle. To actively segregate origins of replication, most prokaryotes rely on the ParAB*S* system (*1, 2*), which consists of a *parS* binding sequence, the ATPase partition protein A (ParA), and the CTPase partition protein B (ParB) (*3–6*). ParB dimers bind to the *parS* sequence, located near the origin of replication, spread laterally and ultimately form ParB-DNA partition complexes that are imperative for DNA segregation (*7–13*). A ParA gradient along the cell axis subsequently segregates these complexes to effectively administer the nascent genomes to the daughter cells (*14, 15*). Proper partitioning of the ParB complexes is vital for bacterial cell survival and has been studied intensively in recent years (*16*), especially since it was recently discovered that ParB proteins from several model organisms utilize CTP to load onto a *parS* sequence (*4, 5, 17*).

A necessary feature for correct chromosome partitioning comes from a particular ability of ParB proteins to clamp the *parS*-DNA as a dimer (*4, 5*) and to subsequently slide along the DNA by diffusion, effectively freeing the *parS* site for new ParB proteins to load. Indeed, ParB has been found to laterally spread over large genomic regions surrounding the *parS* sites *in vivo* (10-15 kb; (*18–22*)), which was reported to be essential for partition complex formation (*9, 10, 16, 18, 19, 23–25*). However, *in vitro* studies showed that single DNA-bound proteins block the diffusion of ParB along DNA very efficiently (*4, 7, 17*), raising the question how spreading can occur in a dense cellular environment, where, with ~1 gene and ~20-50 Nucleoid Associated Proteins (NAPs) per kb (*26, 27*), sliding ParB dimers will continuously run into ‘roadblocks’ that will stall their movement. Indeed, theoretical modelling of a ParB ‘clamping and sliding’ model indicated that such roadblocks dramatically limit the spreading distance on F-plasmids (*28*). Yet, *in vivo* ChIP-seq data do not show strong changes in the ParB occupancy in the vicinity of genes and operons (*18–22, 29*).

Here, we examine the mechanism of ParB spreading using a controlled DNA-stretching assay that allows *in vitro* visualization at the single-molecule level (*30*). First, we verified that CTP and the *parS* site are essential for the loading and diffusion of ParB on DNA. We tethered 42 kb long DNA molecules with a single *parS* site close to the middle (DNA_*parS*_) to a streptavidin-functionalized surface via their 5’-biotin ends (Fig. 1A, S1A). Highly Inclined and Laminated Optical sheet microscopy was used to observe DNA and proteins labelled with different fluorescent dyes (see *Supplementary info,* Fig. S1B-F). ParB was found to exhibit two different types of behaviors, viz., transient non-diffusive binding at various locations (Fig.S2A-C), as well as binding at the *parS* site followed by diffusion (Fig. 1B and S2D-E). Transient binding was short (~5-20 s), independent of the presence of CTP in the medium, and not correlated to the *parS* sequence (Fig 1C, S2B-C). In the presence of both the *parS* sequence and CTP, however, binding initiated predominantly at the location of the *parS* sequence (Fig. 1D and Fig. S2D-H). This binding persisted for an average time of 94 ± 8 seconds (average ± SEM; Fig. 1E), during which time the ParB diffused away from the *parS* site. This residence time is in line with the previously determined CTP hydrolysis rate of 1 CTP per 100 s for ParB_*BSu*_ (*4*), where the hydrolysis of CTP would cause the ParB clamp to destabilize and open, allowing dissociation from the DNA (*4, 5, 19*). Interestingly, we also observed a minor second population of molecules with roughly double the residence time of the first population (188 ± 18 s (average ± SEM), Fig. 1E, S3). This strengthens previous indications that ParB dimers can prolong their residence time on DNA by exchanging CTP molecules after loading (*19*).

**Figure 1.**
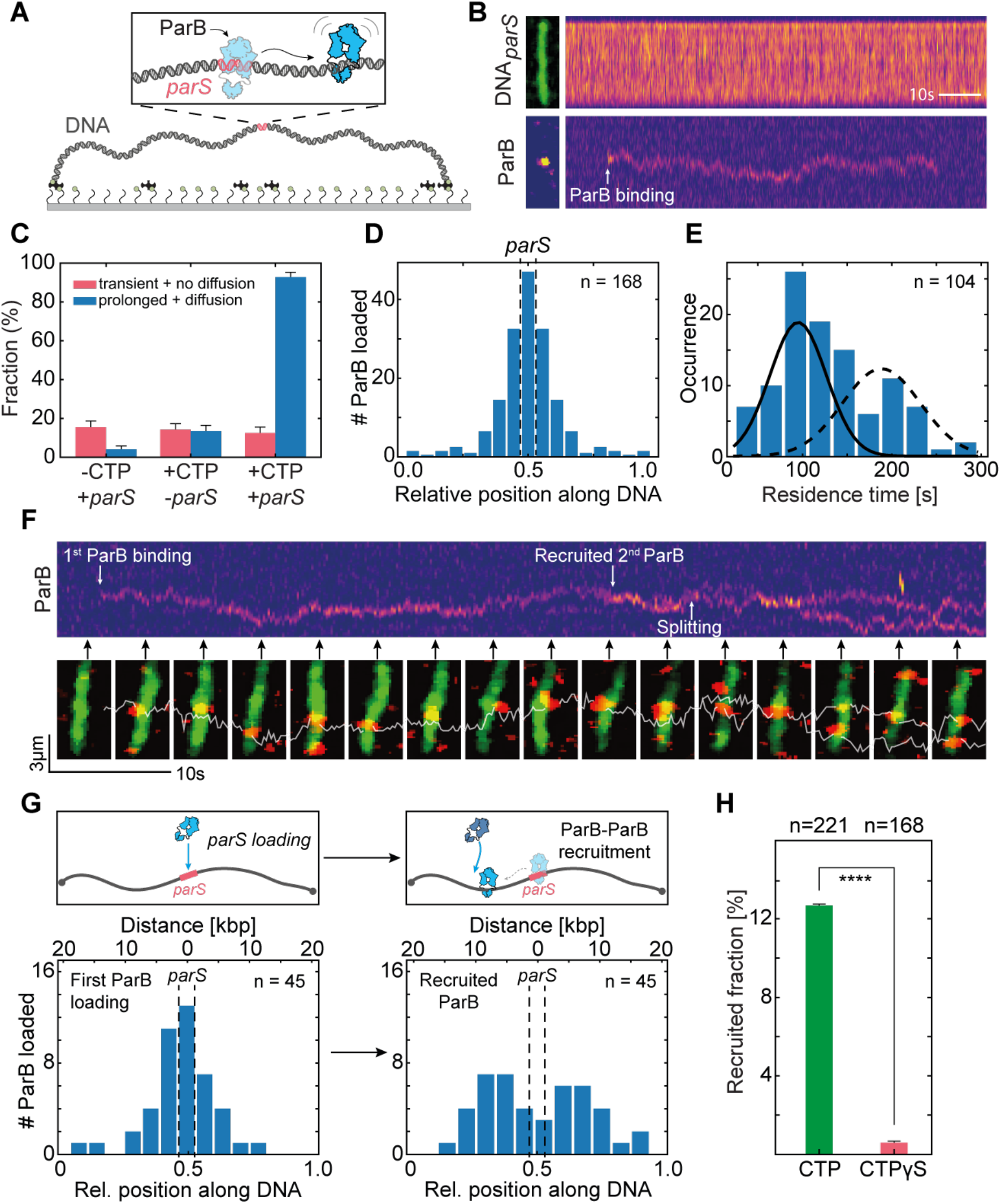
DNA-bound ParB dimers can recruit new ParB dimers independently of *parS.* **A)** Schematic representation of DNA_*parS*_ that is tethered at both its ends to a surface, see also Fig. S1. **B)** Kymographs for DNA_*parS*_ stained with SytoxGreen (top) and ParB^alexa647^ (bottom). Left images show single-frame snapshots of the DNA and ParB at the moment of binding. White arrow indicates ParB loading. C) Fraction of DNA tethers with transient binding with no diffusion and the fraction showing prolonged binding with diffusion of ParB molecules, for various conditions (n = 123, 133, 112 from left to right). Error bars represent binomial proportion confidence intervals. D) Histogram showing loading position of ParB molecules along DNA_parS_. The position of *parS* site is represented by dashed lines, n=168. E) Residence times of diffusing ParB molecules after binding DNA_*parS*_ (n=127). Gaussian fits are represented in black. F) Top: kymograph for ParB^alexa647^. White arrows indicate first ParB-dimer binding, second ParB-dimer binding, and splitting. Bottom: Snapshot images, taken at time points indicated by black arrows. White line indicates the single-particle-tracking trace. G) Histograms of ParB loading positions. Left: First ParB loading at *parS* site. Right: loading position of the recruited second ParB dimer. Top panels are illustrative schemes. H) Recruitment rate in presence of CTP (green) or CTPγS (red). Error bars represent binominal confidence intervals. Statistical significance calculated using chi-squared test for a binomial distribution, χ^2^ (n=389) = 20.196, p<0.00001.

Strikingly, the data also revealed an unexpected behavior that was different from mere loading and diffusion of ParB dimers: a ParB dimer that was previously loaded onto the DNA_*parS*_, was observed to be able to load a new ParB dimer onto the DNA (Fig. 1F, G and S4A-D). This behavior was characterized by the recruitment of a second ParB dimer at the site of an existing one (as evidenced from an increase of the fluorescent intensity), a brief colocalization and conjunct diffusion of the two ParB dimers (for 8.2 ± 3.5 s, average ± SD; Fig. S4D), after which the two dimers split into two independently diffusing ParB dimers (Fig. S4A-C). This intriguing behavior occurred in about 12% of the observed ParB molecules that loaded onto the *parS* site (Fig. 1H). We refer to this type of loading behavior as “ParB-ParB recruitment”. We can distinguish ParB-ParB recruitment from simple subsequent loading of ParB proteins at the *parS* site by evaluating the position along DNA where events occur. As expected, the first ParB dimer loaded at the *parS* site near the middle of the DNA_*parS*_ (Fig. 1F, 1G-left), but the loading of the second, recruited ParB dimer occurred later at a position that was, on average, on 5.2 ± 3.8 kb (average ± SD, n=46) away from the *parS* site (Fig. 1F, 1G-right). Accordingly, the first ParB loaded at *parS* site and already diffused over a significant distance before recruiting a new ParB dimer at a distant location (Fig. 1G-right). Interestingly, after ParB-ParB recruitment, we observed that *both* ParB dimers resided on the DNA for ~72 s, i.e., for a similar amount of time as single diffusing ParB before recruitment (~66 s, Fig. S4C). The recruitment thus approximately doubled the residence time of the initially loaded ParB dimer to ~138 s (Fig. S4C). Notably, the ParB-ParB recruitment process was highly dependent on the presence of CTP, and occurred hardly at all in the presence of the slowly hydrolysable nucleotide variant CTPγS (Fig. 1H)(*25*). Hereafter, we refer to these recruitment events where the second ParB loaded at a DNA position adjacent to the first one, as *‘in-cis’* (Fig. 2A-left).

**Figure 2.**
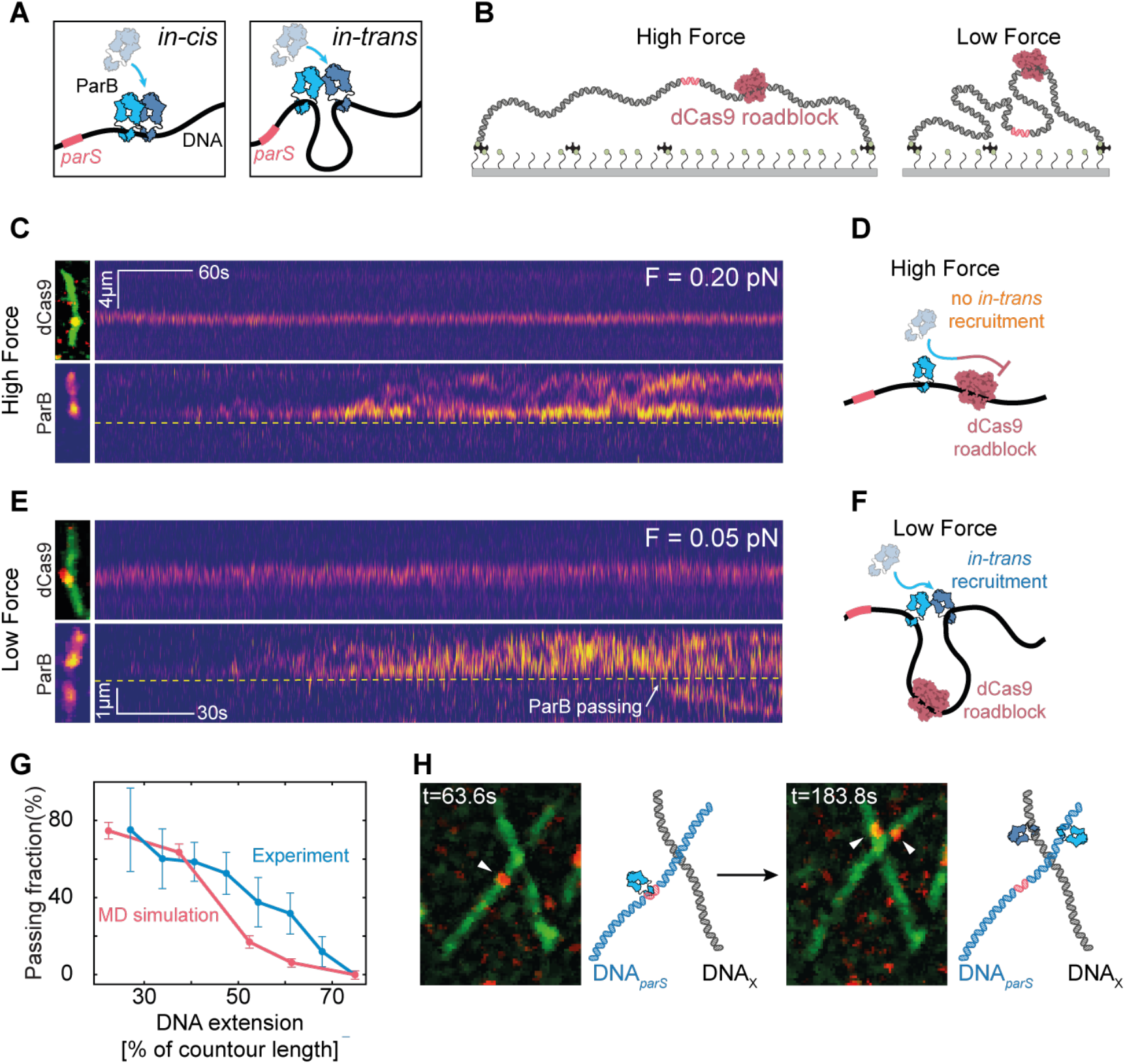
ParB can bypass roadblocks by *in-trans* ParB-ParB recruitment. **A)** Illustration of *in-cis* (left) and *in-trans* (right) recruitment events. **B)** Schematic of the roadblock experiment for a large (left, high force) and small DNA_*parS*_ end-to-end distance (right, low force). **C)** Kymograph for dCas9^alexa549^ (top), and ParB^alexa647^ (bottom) for DNA_*parS*_ at 0.20 pN. Snapshots of dCas9^alexa549/SytoxGreen^DNA*_parS_* overlay and ParB^alexa647^ signal are provided on the left. Yellow dashed line indicates the position of the dCas9^alexa549^. **D)** Cartoon of the DNA_*parS*_ at high force where ParB molecules cannot be recruited across the roadblock. **E)** Same as panel C but data at a force of 0.05 pN. White arrow denotes a passing event. **F)** Same as panel D but for low force where ParB molecules can be recruited *in-trans* across the dCas9–roadblock. **G)** Fraction of DNA molecules that exhibit ParB dimers passing the dCas9-roadblock, displayed versus stretching length of the DNA. Error bars represent binomial proportion confidence intervals (blue, n = 122), and standard deviation from simulation cycles (pink, n=128). **H)** Left: Snapshot of the DNA_*parS*_-DNA_X_ crossing at the time of the first ParB binding to *parS* site, t=105s. Right: Snapshot of the same field of view at time t=306s, where a second ParB was recruited onto the DNA_X_ by *in trans* recruitment. White arrows indicate the positions of ParB dimers. Cartoon representations are provided on the right of the snapshot frames.

By contrast, we observed that ParB-ParB recruitment events can also happen *‘in-trans’*, where the second ParB got loaded onto a DNA position that is genomically distant, but transiently proximal in 3D space via DNA looping (Fig. 2A-right). Since these types of *in-trans* events could potentially allow for the passage of DNA-bound roadblocks in the crowded *in vivo* environment, we designed an experiment to test for *in-trans* recruitment in the presence of DNA-bound roadblocks. We placed a firmly bound dCas9 protein on one side of the *parS* sequence on the DNA (Fig. 2B) and tuned the end-to-end distance of the DNA to 65% of its total contour length (corresponding to a stretching force of 0.20 pN within the DNA (*31*)). In these conditions, we observed that the dCas9 roadblock efficiently prevented the diffusion of ParB proteins (Fig. 2C, D and Fig. S5A-E), in line with findings with other DNA-bound roadblocks (*4, 7, 17*). However, when repeating the same experiment for a lower end-to-end distance (27% of the DNA*_parS_* contour length; *F =* 0.05 pN (*31*)), we observed events where ParB proteins would cross the roadblock and continue one-dimensional diffusion along the DNA on the other side of dCas9 (Fig. 2E, F and S6A-E, Movie S1). We attribute this behavior to *in-trans* ParB-ParB recruitment. The bypassing of the Cas9 roadblock was also visibly apparent in the cumulated ParB fluorescence intensity signal, that rapidly increased on the *parS* side, while the non-*parS* side displayed discrete increases in intensity only for the non-stretched DNA_*parS*_ (Fig. S5C, E and S6B, D).

The tension within the DNA appeared to have a significant effect on the success rate of crossing the roadblock. As Fig. 2G displays, the fraction of DNA molecules where ParB successfully passed the dCas9 roadblock decreased from ~80% at 0.05 pN (27% of the total DNA contour length) to zero at 0.35 pN (75% contour length (*31*)). These data are consistent with *in-trans* ParB-ParB recruitment, where at low stretching forces, regions that are distant in DNA sequence can come into physical contact through bending and looping of the DNA via thermal fluctuations. We further corroborated these findings by observing the ParB-mediated *in-trans* loading of a new ParB dimer onto a *different* DNA molecule that was spatially nearby, where this DNA did not contain an endogenous *parS* site (Fig. 2H, S7). Here, as previously, we bound the DNA_*parS*_ to the surface but then subsequently bound a new DNA molecule lacking the *parS* site (DNA_X_), perpendicular to the originally bound DNA_*parS*_ (Fig. S7A-B). In this assay, we observed events where ParB first specifically bound the DNA_*parS*_, then reached the crossing point by random diffusion, where it recruited a new ParB molecule onto the DNA_X_ (Fig. S7C, dashed line), yielding a transfer and subsequent independent one-dimensional diffusion of the newly recruited ParB dimer on the DNA_X_ molecule (Fig. 2H, S7C).

Using coarse-grained molecular dynamics simulations, we modeled the passage of DNA-bound roadblocks by *in-trans* binding, and found a similar strong force dependence of the passing fraction of roadblocks. We simulated DNA tethers containing a blocking particle which strictly prohibited ParB diffusion through it (Fig. 3A), whereas concurrently ParB could exhibit *in-cis* or *in-trans* recruitment with an inbuilt rate (Fig. 3B, see *Methods*). While in the absence of a roadblock, both *in-cis* and *in-trans* recruitment increased the lateral spreading of ParB on the DNA (Fig. S8A), the presence of a roadblock allowed ParB to spread beyond it only through *in-trans* recruitment (Fig. 3C-D, S8B-D). Upon quantifying the passing fractions at different DNA end-to-end lengths and *cis/trans* recruitment ratios, we obtained the closest comparison with the experimental data at *cis/trans* ratio of ~0.1 (Fig. 2G, 3E). The modeling at very low forces verified the notion that contacts between regions on opposing sides of the roadblock occur sufficiently often to allow for ParB to overcome roadblocks via ParB-ParB recruitment.

**Figure 3.**
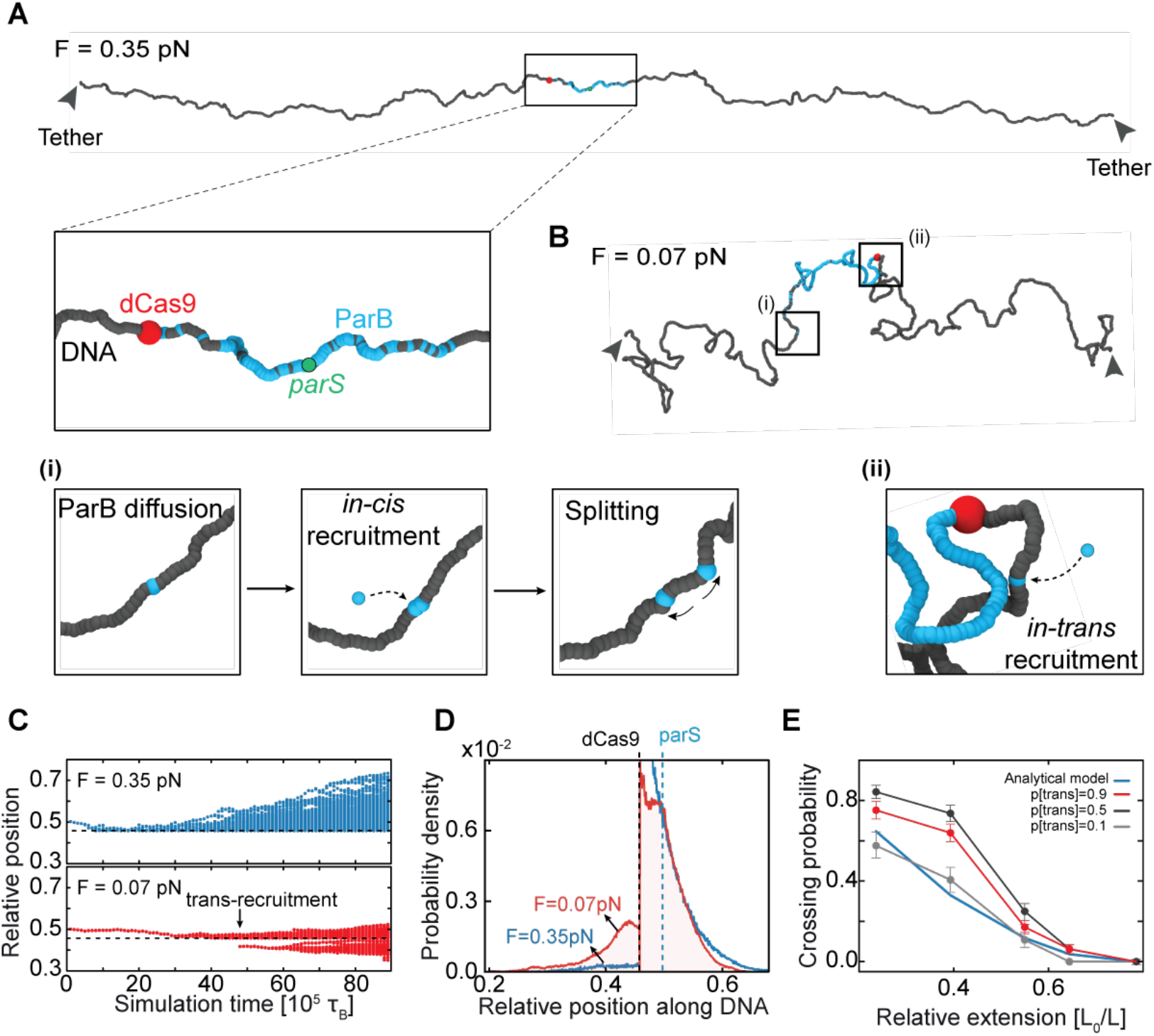
DNA looping via thermal fluctuations allows *in-trans* ParB-ParB recruitment and spreading across roadblocks. **A)** Single frame snapshot of the DNA tethers in the MD simulations. DNA tether end-to-end length 11*μ*m (i.e., 75% contour length, force of 0.35pN). Zoomed region highlights the region proximal to *parS* site. Tether has a *parS* site in the middle (green), and simulated dCas9-roadblock on one side (orange). ParB represented in yellow. **B)** Single frame snapshot of low-force DNA molecule in MD simulations. DNA tether end-to-end length 5.5*μ*m (i.e., 35% contour length, force of 0.07pN). Highlighted regions (a) and (b) show *in-cis* and *in-trans* recruitment events, respectively. Color-coding same as in panel A. **C)** Simulated kymographs representing distributions of ParB positions during the simulation. Top: kymograph for DNA tether at 0.35pN; bottom: same at F=0.07pN. Black arrow indicates *in-trans* recruitment event. **D)** Histogram of cumulated ParB probability density from MD simulations at different tether lengths (blue 11*μ*m, red 5.5*μ*m) averaged over n=64 simulations. Dashed line indicates the position of dCas9 roadblock. **E)** Fraction of simulated DNA tethers that exhibit ParB dimers passing the dCas9-roadblock at different *cis/trans* ratios. The theoretical passing rate (integration of polymer looping probability, see Fig. S8E) is shown in blue. Error bars represent standard deviation from simulation cycles (n=128).

We found CTP hydrolysis to be crucial for the occurrence of ParB-ParB recruitment (Fig. 1H, S9, Movie S2). To examine this further, we replaced CTP by the poorly hydrolysable CTPγS (Fig. S9A), which allows for the initial loading (clamp closing) but reduces opening of the ParB clamp upon hydrolysis (Fig. S9B-E)(*4, 7, 19, 25*). This resulted in a low and length-independent passing fraction, indicating an essential role of CTP hydrolysis in the *in-trans* ParB-ParB recruitment (Fig. S9F-H). As CTP hydrolysis is tightly connected with ParB clamp re-opening and dissociation from the DNA, it is likely that the N-terminal domain of ParB plays a role in the recruitment process. Interestingly, ParB mutants defective in CTP hydrolysis previously showed an altered distribution around *parS* in *B. subtilis,* with a reduced spreading against the orientation of highly expressed rRNA genes and the DNA replication machinery, and an increased spreading in the opposite direction (*19*). We suggest that this distribution for the mutant ParB arises from one-dimensional diffusion only (equivalent to the clamping and spreading model), while the spreading of the wild-type protein additionally relies on ParB-ParB recruitment that can overcome head-on encounters with protein machineries such as RNA polymerases or replication forks. Similar observations were made for ParB in *C. crescentus* (*25*), whereas by contrast, *Myxococcus* ParB mutants showed extensive spreading in both directions(*32*), possibly by unhindered 1D diffusion due to the differences in life cycles (i.e., lower gene transcription and replication rates – see *33, 34*).

Based on our findings, we propose a model for ParB spreading (Fig. 4) that expands the well-known clamping and sliding model. First, ParB loads onto the *parS* site and spreads away by one-dimensional diffusion (Fig. 4i), until it hydrolyzes the bound CTP. In our model, however, unlike commonly assumed, the hydrolysis does not necessarily imply the dissociation of ParB from the DNA. Instead, we hypothesize that the ParB dimer may briefly reside in an intermediary state after hydrolyzing CTP, where one of the monomers in not fully engaged in N-terminal dimerization (*19*). Given abundant CTP in the surrounding buffer, the CDP can now exchange to a new CTP molecule, which leaves the CTP-bound free N-terminal part of the protein with two possibilities: (1) either re-connect with the N-terminus on the adjacent monomer of the ParB to re-close the clamp and continue diffusion (Fig. 1E, 4ii) (*19*), or (2) connect to the N-terminus of a different ParB dimer that is nearby in the solution. The second scenario can lead to recruitment of a second ParB dimer to the nearby DNA, which can occur either *in-cis* or *in-trans* (Fig. 4iii). *In-trans* recruitment, in this fashion, can consequently result in DNA-roadblock passing and free diffusion on the other side. The precise molecular mechanism of how the recruited ParB releases the dimer-dimer connection, resulting in separate independent diffusion of both ParB dimers, remains unknown (Fig. 4iii and 4iv).

**Figure 4.**
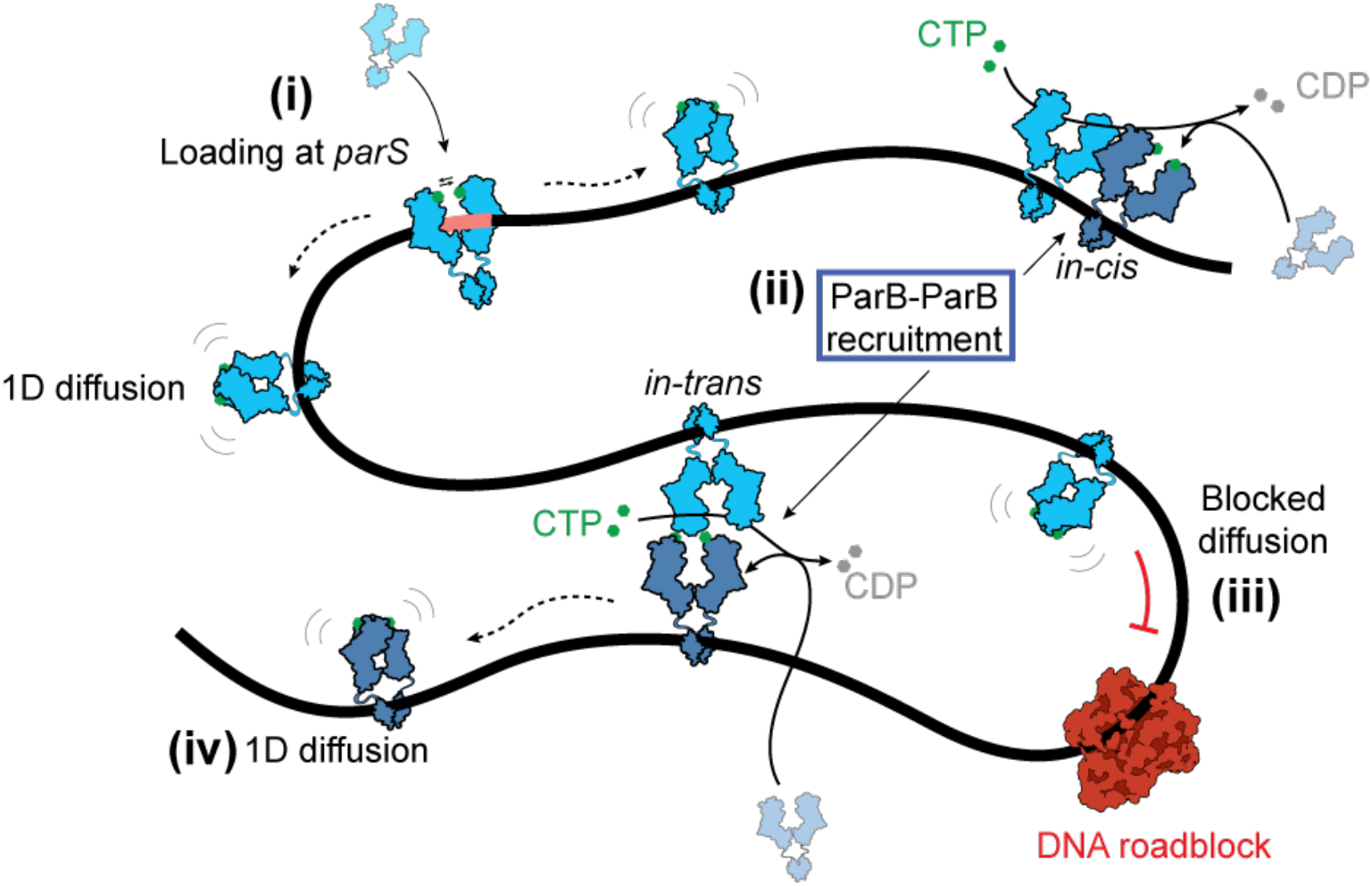
Model of ParB spreading and recruitment. Cartoon of the various ParB modalities on DNA: (i) ParB dimer loads at the *parS* site, dimerizes upon closing its N-termini, losing its affinity to the *parS*-site, and diffusing away as a closed clamp. Subsequently (ii), ParB hydrolyzes CTP and enters an intermediary state where it can bind and recruit another ParB dimer to a nearby DNA, which can occur *in-cis* or *in-trans.* By exchanging CDP and Pi after hydrolysis for a new CTP molecule from solution, ParB dimer can also re-form a clamped structure and continue diffusing. While one-dimensional diffusion is blocked by DNA-roadblocks (iii), ParB dimers that are recruited *in-trans* across the roadblock can continue diffusion along the DNA, yielding an expanded spreading of ParB along DNA.

Overall, *in-trans* ParB-ParB recruitment is consistent with the extensive spreading of ParB along genomic DNA that contains many roadblock proteins, Furthermore, it offers an alternative mechanism to the ‘stochastic binding’ that was proposed in previous models (*28*) to explain the presence of ParB at large distances from *parS* sites in *in vivo* ChIP-seq data. We speculate that ideal hotspots for ParB *in-trans* recruitment events may additionally be found in regions that are brought into close proximity of a *parS* site by the action of an SMC complex (*35, 36*). Indeed, recent reports on Hi-C maps detected in strains that were constructed for testing SMC collisions *in vivo*, showed the co-alignment of DNA flanking a *parS-free* region which is characteristically only found at *parS* sites. Physical contact of DNA with a genomically-distant *parS*-containing region may lead to *in-trans* ParB-ParB recruitment and create *parS-free* ParB-DNA structures suitable for further off-site SMC recruitment (*37*).

Our findings uncover a new pathway for ParB proteins to cooperatively cover large distances on DNA that is loaded with DNA-binding proteins. It involves the combination of lateral 1D diffusion along the DNA and a new type of CTP-hydrolysis-dependent ParB loading (ParB-ParB recruitment) which can occur irrespective of the *parS* site. Both our experimental and simulation data showed that *in-trans* ParB-ParB recruitment can account for overcoming DNA-bound roadblocks at low forces where DNA forms a fluctuating polymer blob that facilitates frequent DNA-DNA encounters. Since both the ParB concentration and the DNA-contact frequency are higher in the tightly packed genome within a bacterial cell compared to our single-molecule experiments, we expect ParB-ParB recruitment to be a common mechanism *in vivo*, where it may facilitate the collective spreading distance of ParB proteins on the protein-bound DNA.

## Acknowledgments

We thank Alejandro Martin Gonzalez, Eugene Kim, Florian Patrick Bock and Richard Janissen for useful discussions, and Richard Janissen for sharing software for force estimation on the DNA tether based on the end-to-end distance.

## Author contributions

Conceptualization: MT, SG, CD. Experiments: MT. Formal analysis: MT, BP, RB. Methodology: MT, JvdT, CD, MP, DM. Protein purification: HA, YMS. Software: BP. Simulations: MP, DM. Visualization: MT. Funding acquisition: DM, SG, CD. Supervision: DM, SG, CD. Writing – original draft: MT, CD. Writing – review & editing: MT, HA, YMS, MP, RB, BP, JvdT, DM, SG, CD.

## Data and materials availability

All data are available in the main text or the supplementary material.

