## Supplementary materials for "*ParS*-independent recruitment of the bacterial chromosome-partitioning protein ParB"

### Supplementary material

#### Materials and Methods

*ParB purification and fluorescent labelling*

We prepared ParB expression constructs using pET-28a derived plasmids through Golden-gate cloning. We expressed untagged recombinant proteins in *E. coli* BL21-Gold (DE3) for 24 h in ZYM-5052 autoinduction medium at 24°C. Purification of $ParB^{L5C}$ variants was performed as described before(*1*). Briefly, we pelleted the cells by centrifugation and subjected them to lysis by sonication in buffer A (1 mM EDTA pH 8, 500 mM NaCl, 50 mM Tris-HCl (pH 7.5), 5 mM β-mercaptoethanol, 5 % (v/v) glycerol, and protease inhibitor cocktail (PIC, SigmaAldrich). We then added ammonium sulfate to the supernatant to 40% (w/v) saturation and kept stirring at 4°C for 30 min. We centrifuged the sample and collected the supernatant, and subsequently added ammonium sulfate to 50% (w/v) saturation and kept stirring at 4°C for 30 min. We collected the pellet (containing ParB) and dissolved it in buffer B (50 mM Tris-HCl (pH 7.5), 1 mM EDTA pH 8 and 2 mM β-mercaptoethanol). The sample was also diluted with buffer B to achieve a conductivity of 18 mS/cm before loading onto a Heparin column (GE healthcare). We used a linear gradient of buffer B containing 1 M NaCl to elute the protein. After collecting the peak fractions, we diluted them in buffer B to 18 mS/cm conductivity, and loaded onto HiTrap SP columns (GE healthcare). For elution, we used a linear gradient of buffer B containing 1 M NaCl. Collected peak fractions were loaded directly onto a Superdex 200-16/600 pg column (GE healthcare) preequilibrated in 300 mM NaCl, 50 mM Tris-HCl (pH 7.5), and 1 mM TCEP. For fluorescent labeling, we incubated purified $ParB^{L5C}$ variant with either TMR-maleimide, Alexa647-$C_2$-malemeide, or JanielaFluor646-malemeide at a 1:1.2, 1:10, and 1:5 (protein:dye) molar ratio, respectively. We incubated the mixture for 15 min on ice, centrifuged for 10 min and then eluted from a spin desalting column (Zeba) and flash frozen in liquid nitrogen.

*Labelled ParBs CTP-hydrolysis assays (alexa488, TMR, alexa647)*

We test for the activity of the labelled-$ParB^{L5C}$ by its ability to hydrolyze CTP in the presence of $DNA_{parS}$. For this reason, we measured the hydrolysis rate by Malachite Green colorimetric detection. In brief, we prepared a mixture of 2x[CTP] and 2x 40bp $DNA_{parS}$ concentration in reaction buffer (150 mM NaCl, 50 mM Tris-HCl (pH 7.5), 5 mM $MgCl_2$) and placed it on ice. We added an equal volume of 2x solution, containing the purified labelled-$ParB^{L5C}$, protein to the 2xCTP/DNA mixture and left to incubate at 25°C for 1 h in a PCR machine. In parallel, we prepared phosphate blanks. The samples were then diluted 4-fold with MiliQ water, and subsequently mixed with 20 µl working reagent (SigmaAldrich) and transferred to a flat bottom 96-well plate. The plate was left to incubate at 25°C for 30 min, and we measured the absorbance at a wavelength of 620 nm. We used the absorbance values from the phosphate standard samples to plot an $OD_{620}$ versus phosphate concentration standard curve. Using the standard curve, we converted raw values to rate values, and calculate the absolute rates by normalizing for protein concentration.

*Construction and purification of 42kb $DNA_{parS}$ construct*

For the construction of a long linear $DNA_{parS}$, we used a large 42kb cosmid-i95 reported previously (*2*) and a synthetic construct containing the *parS* site (Integrated DNA Technologies, Table. S1, underlined sequence). First, we linearized the i95 cosmid using the PsiI-v2 restriction enzyme (New England Biolabs). Next, we dephosphorylated the remaining 5'-phosphate groups using Calf Intestinal Alkaline Phosphatase for 10 min at 37°C, followed by heat inactivation for 20 min at 80°C (Quick CIP, New England Biolabs). We added the 5'-phospho group on the synthetic *parS* fragment by adding a T4 kinase for 30 min at 37°C and heat-inactivated 20 min 65°C in 1x PNK buffer supplemented with 1 mM ATP (T4 PNK, New England Biolabs). Next, we ligated the two fragments together using a T4 DNA ligase in T4 ligase buffer (New England Biolabs), containing 1 mM ATP overnight at 16°C. The final cosmid construct was transformed into *E. coli* NEB10beta cells (New England Biolabs), and we verified the presence of insert by sequencing using JT138 and JT139 (Table S1). To prepare a linear fragment adapted for flow cell experiments, we isolated cosmid-i95 via a midiprep using a Qiafilater plasmid midi kit (Qiagen). The cosmid-i95 was then digested for 2 h at 37°C and heat-inactivated for 20 min at 80°C using AjuI restriction enzyme (ThermoFischer Scientific). Linear DNA constructs for three-color experiments with roadblocks were constructed in the same way using the SpeI-HF restriction enzyme (New England Biolabs).

Next, we constructed the 5'-biotin handles by a PCR from a pBluescript SK+ (Stratagene) using 5'-biotin primers JT337 and JT338 (Table S1), to get a final 1245 bp fragment. The PCR fragment was digested using the same procedure described for cosmid-i95, resulting in ~600 bp 5'-biotin-handles. Finally, we mixed the digested cosmid-i95 and handles in a 1:10 molar ratio and ligated them together using T4 DNA ligase in T4 ligase buffer (New England Biolabs) at 16°C overnight, which was subsequently heat-inactivated for 25 min at 65°C. We cleaned up the resulting linear (42 +1.2) kb $DNA_{parS}$ construct from the access handles using an ÄKTA Pure (Cytiva), with a homemade gel filtration column containing 46 ml of Sephacryl S-1000 SF gel filtration media, run with TE + 150 mM $NaCl_2$ buffer at 0.2ml/min. We stored the collected fractions as aliquots after snap-freezing them by submerging them in liquid nitrogen.

*Preparation and binding dCas9 roadblock to $DNA_{parS}$*

To form the dCas9-alexa549 roadblock complex, we initially prepared a tr-crRNA duplex by mixing universal 67mer trRNA and a custom-designed crRNA (Table S1), in a duplex buffer (Integrated DNA Technologies), to a final concentration of 10 µM each. The mixture was incubated at 95°C for 5 min, and then slowly cooled to 4°C by decreasing the temperature for 5°C every 5 min over the course of 1.5 hours. We next incubated the tr-crRNA solution with the dCas9 in the "binding buffer" (2 µM tr-crRNA complex, 1 µM dCas9-SNAP (New England Biolabs), 1x NEB3.1 buffer) for 10 min at 37°C, before placing it on ice. Following the tr-crRNA-dCas9 complex formation, we bound it to the $DNA_{parS}$ by incubating the $DNA_{parS}$:tr-crRNA-dCas9 in molar ratio 1:50 for 60 min at 37°C. We labelled the dCas9-SNAP by adding alexa546-BG to a final concentration of 1 µM for 30 min at room temperature before flowing it into the flow cell.

*Single-molecule visualization assay*

The surface of imaging coverslips was prepared as previously described (*3*), with the addition of surfaces being pegylated 5x24h. For immobilization of 42kb $DNA_{parS}$, we introduced 50 µl of ~1 pM of 5'-biotinylated-$DNA_{parS}$ molecules at a flow rate of 3 – 14 µl/min, depending on the desired end-to-end length in the experiment, in T20 buffer (40 mM Tris-HCl (pH 8.0), 20 mM NaCl, 25 nM SytoxGreen (SxG, ThermoFisher Scientific)). Immediately after the flow, we further flowed 100 µl of the wash buffer (40 mM Tris–HCl, pH 8.0, 20 mM NaCl, 65mM KCl, 25 nM

SxG) at the same flow rate to ensure stretching and tethering of the other end of the DNA to the surface. By adjusting the flow, we obtained a stretch of around 25−80% of the contour length of DNA. Next, we flowed in the imaging buffer (40 mM Tris-HCl, 2 mM Trolox, 1 mM TCEP, 10 nM Catalase, 18.75 nM Glucose Oxidase, 30 mM Glucose, 2.5 mM $MgCl_2$, 65 mM KCl, 0.25 µg/ml BSA, 1 mM CTP, 25 nM SxG) without ParB protein at the same flow rate to maintain identical conditions before and after protein addition. Real-time observation of ParB diffusion was carried out by introducing ParB (0.1-1 nM) in the imaging buffer. We used a home-built objective-TIRF microscope to achieve fluorescence imaging. We used alternating excitation of 488-nm and 646-nm, 561-nm lasers in Highly Inclined and Laminated Optical sheet (HiLo) microscopy mode, to image SxG-stained DNA and Alexa647- or TMR-labelled ParB using. When imaging Alexa488-ParB, we used continuous 488 nm excitation in the absence of SxG. All images were acquired with an PrimeBSI sCMOS camera at an exposure time of 100 ms for dual-color experiments and 60 ms for three-color experiments, with a 60x oil immersion, 1.49NA CFI APO TIRF (Nikon).

*Crossed $DNA_{parS}$-$DNA_X$ assay*

The surface was prepared, and binding of the $DNA_{parS}$ was done the same as described above. The flow cell was then blocked at the primary inlet (Fig. S7) and resumed from a secondary inlet to establish a cross flow that was oriented under a substantial angle with the flow that stretched $DNA_{parS}$ on the surface. In this second flush, DNA that contained no *parS* site was stretched onto the surface, until sufficient number of crossed $DNA_{parS}$-$DNA_X$ events was observed. We performed the remainder of the experiment identically to the previously described protocol with a fixed concentration of 500 pM ParB.

*Data analysis of single-molecule imaging traces*

We resolved the initial loading positions of ParB by calculating the mean pixel position over the first full second of the ParB traces from the kymograph, and then determining that value relative to the DNA ends, i.e.,

$$Relative\ loading\ position = \frac{(x_{end} - x_{load\ avg.})}{(x_{end} - x_{start})}.$$

All loading positions obtained like this were pulled together and represented in a histogram using a custom-written python script (Fig. 1D, S2B, H, S9). We determined the residence time of ParB molecules by measuring the length of traces of single ParB proteins by their fluorescence (Fig. 1E, S3A, S4C, D, S9E). We fitted the histogram with multiple Gaussian populations using the *pomegranate* module (*4*) and computed the Bayesian Information Criterion (BIC, Fig. S3B-top) and Akaike Information Criterion (AIC, Fig. S3B-bottom) for each fit in order to determine the optimal number of populations underlying the data. The mean and standard error of the mean were obtained via bootstrapping of all data ($n_{iterations}$ = 5000, Fig 1E). Similarly, we obtained the ParB residence time before and after recruitment (Fig. S4C, D).

We obtained the *parS* and non-*parS* arm intensities in dCas9-roadblock experiments (Cf. Fig. 2C, E) by selecting the corresponding regions in the kymograph before (~100 frames) and after (~2000 frames) the observable binding of the first ParB molecule to the DNA. Pixel values before the initial binding, representing noise, were averaged over the 100 frames and subtracted from each frame in the region after the initial molecule bound. The roadblock position was determined as the maximum value over the time-averaged kymograph in the 561 nm channel after applying the Savitzky–Golay filter implemented from the *scipy* python module, with a window length of 11 frames, and order 1. ParB signal intensity was obtained by applying a median filter the kymograph from 647 nm channel with a kernel size of 21 frames to account for the signal noise and the noise from the wiggling of the DNA molecule. We discarded intensity data from a window of 5 pixels above and below the dCas9 position to account for "leaking" fluorescence due to fixed pixel position of dCas9 (at the peak of Savitzky–Golay curve) and simultaneous wiggling DNA and ParB signal with it. The raw data was plotted on the same graph as the median filter curves, in the background.

To determine the fluorescence intensity of ParB proteins before, during, and after recruitment events, single discernable ParB proteins were tracked on kymographs as described above. We then integrated the fluorescence from the three surrounding pixels of the track position for 20 frames before, during and after the recruitment event. For visualization of example recruitment events (Fig. S4B) we temporally averaged 20 frames of kymographs to obtain a profile across the DNA length and identified one or two peaks (before/during and after recruitment, respectively) by calling a Gaussian Mixture Model from the python *sklearn* module with the respective number of components.

*Modelling and molecular dynamics simulations*

We modelled the DNA as a semiflexible polymer made of $1400$ spherical beads of size $\sigma = 5.5\ nm = 16bp$. We placed *parS* in the middle of the chain where ParB beads were recruited and subsequently diffuse away. In addition, on 2/3 of the DNA chain, we placed a roadblock representing the dCas9 enzyme. The beads interact purely by excluded volume following the shifted and truncated Lennard-Jones (LJ) force field

$$U_{LJ}(r,\sigma) = \begin{cases} 4\varepsilon\left[\left(\frac{\sigma}{r}\right)^{12} - \left(\frac{\sigma}{r}\right)^{6} + \frac{1}{4}\right] & for\ r \le r_c \\ 0\ otherwise \end{cases} \quad , \quad (1)$$

where $r$ denotes the distance between any two beads and $r_c = 2^{\frac{1}{6}}\sigma$ is the cut-off. We defined the bonds between two monomers along the DNA contour length by the finite extensible nonlinear elastic (FENE) potential, given by

$$U_{FENE}(r) = -0.5kR_o^2 \log\left(1 - \left(\frac{r}{R_o}\right)^2\right)\ for\ r \le R_o \quad , \quad (2)$$

with $k = 30\varepsilon/\sigma^2$ the spring constant, $\varepsilon$ is thermal energy, and $R_o = 1.6\sigma$ the maximum length of the bond. We introduced the persistence length of the DNA chain as a bending potential energy between three consecutive beads given by

$$U_{bend}(\theta) = k_\theta(1 - cos\theta) \quad , \quad (3)$$

where $\theta$ is the angle between two bonds and $k_\theta = 10\ k_BT$ is the bending stiffness constant, corresponding to a persistence length of about 10 beads or ~55 nm. We modelled ParB in this system by calling, within the LAMMPS engine, an external program that modifies the types of the beads. At $t = 0$, we loaded a ParB protein onto the *parS* site and allowed to diffuse with a constant

$D = 0.05\frac{\mu m^2}{s} = 0.05\left(\frac{182^2\sigma^2}{2.8\ 10^8 dt}\right) = 0.062\frac{\sigma^2}{10^4 dt}$. This is done by updating the position of a loaded ParB protein either to the left or to the right with probability 0.125 every $10^4 dt = 10^2\tau\_B$ timesteps (recall that MSD = 2 Dt in a 1D system, hence why the jump probability is twice the diffusion coefficient D). The diffusion cannot happen (the move is rejected) if the attempt brings a ParB protein either on top of another ParB or on top of dCas9.

On top of diffusion, we added a recruitment process at a rate of $10^{-6}\tau_B^{-1}$, i.e., on average every $10^6\tau_B$ timesteps (0.35 seconds) another ParB was recruited by a loaded ParB. When this happened, the recruitment could stochastically happen *in-cis* (with probability $p_c$) or *in-trans* (with probability $p_T = 1 - p_C$). If the former was selected, one of the two adjacent beads was selected at random and, if unoccupied and not the dCas9 bead, a new ParB was added onto the chain. Otherwise, if the latter '*in-trans*' mechanism was selected, we computed the list of 3D proximal neighbors which should be (i) within an Euclidean cutoff distance and (ii) farther than the second-nearest neighbor in 1D (i.e. the first and second nearest neighbors cannot be picked). The 3D Euclidean cut-off was set to $2\,\sigma = 11\,nm$ and because of this, the first nearest neighbour was always within 3D reach. To avoid this artificial contribution, we neglected the first two nearest neighbours for trans recruitment. The theoretical bypass probability was given by the integral of the looping probability (Fig. S8E) as

$$P_{bypass} \propto \int_2^\infty P_{loop}(x)dx \ .$$

Once the list of 3D neighbors was compiled, we randomly picked one of these from the list (if not empty), loaded a new ParB protein and resumed the Langevin simulation.

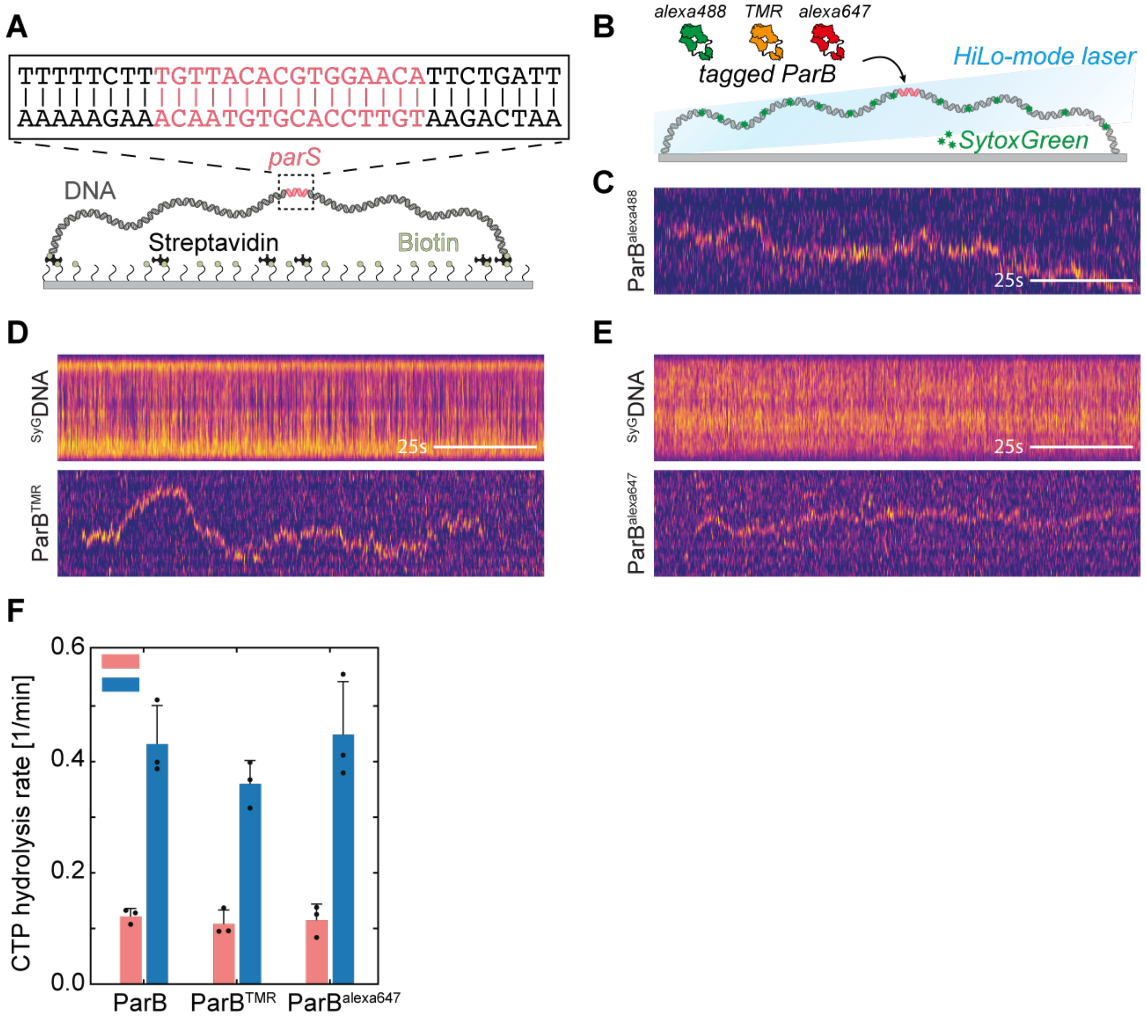


**Figure S1. ParB exhibits one-dimensional diffusion irrespective of the fluorescent tag. A)** Schematic representation of the DNA$_{parS}$ that is tethered at both its ends to a surface. Zoomed region shows parS sequence. **B)** Schematic representation of HiLo imaging setup using different fluorescent tags on ParB as well as SytoxGreen for DNA$_{parS}$ staining. **C)** Kymograph showing ParB diffusion using ParB$^{alexa488}$. Scale bar = 25s. **D)** and **E)** Kymographs for DNA$_{parS}$ stained with SytoxGreen (top) and ParB$^{TMR}$ (bottom) or ParB$^{alexa647}$ (bottom), respectively. Scale bar = 25s. **F)** CTP hydrolysis assay with Malachite Green for unlabelled, TMR- and Alexa647-labelled ParB proteins. The rates were normalized to the blank phosphate controls (see Methods). Error bars represent standard deviation from a triplicate (presented as black circles).

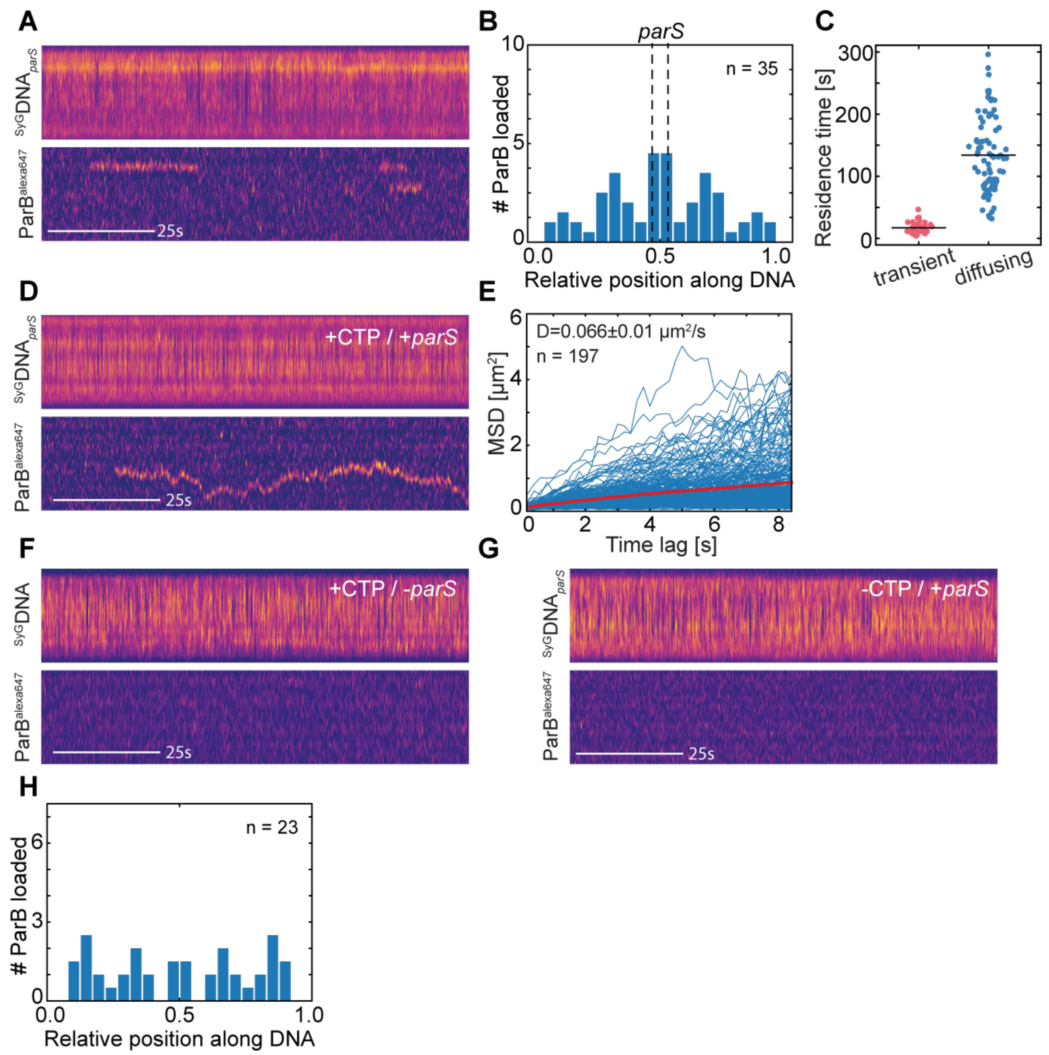


**Figure S2. ParB spreading by diffusion occurs in the presence of *parS* sequence and CTP. A)** Kymographs for DNA$_{parS}$ stained with SytoxGreen (top) and ParB$^{alexa647}$ (bottom). **B)** Mirrored histogram representing transient binding position of ParB molecules relative to the DNA$_{parS}$ ends. The position of *parS* site is represented by dashed lines. n=35. **C)** Residence times of transiently bound non-diffusing (pink) and *parS*-bound and diffusing ParB dimers (blue). **D)** Kymographs for DNA$_{parS}$ stained with SytoxGreen (top) and ParB-alexa647 (bottom) in the presence of both CTP and *parS* site. **E)** Mean square displacement of the diffusing ParB molecules loaded at *parS* site. Apparent diffusion coefficient is on average D= 0.066±0.01 μm$^2$/s, n= 197. **F)** Kymographs for DNA$_{parS}$ stained with SytoxGreen (top) and ParB$^{alexa647}$ (bottom) in the absence of *parS*-site. **G)** Idem in the absence of CTP. **H)** Mirrored histogram for the binding position on non-specifically bound ParB dimers (+CTP/-*parS*) relative to the DNA$_{parS}$ ends. n=23.

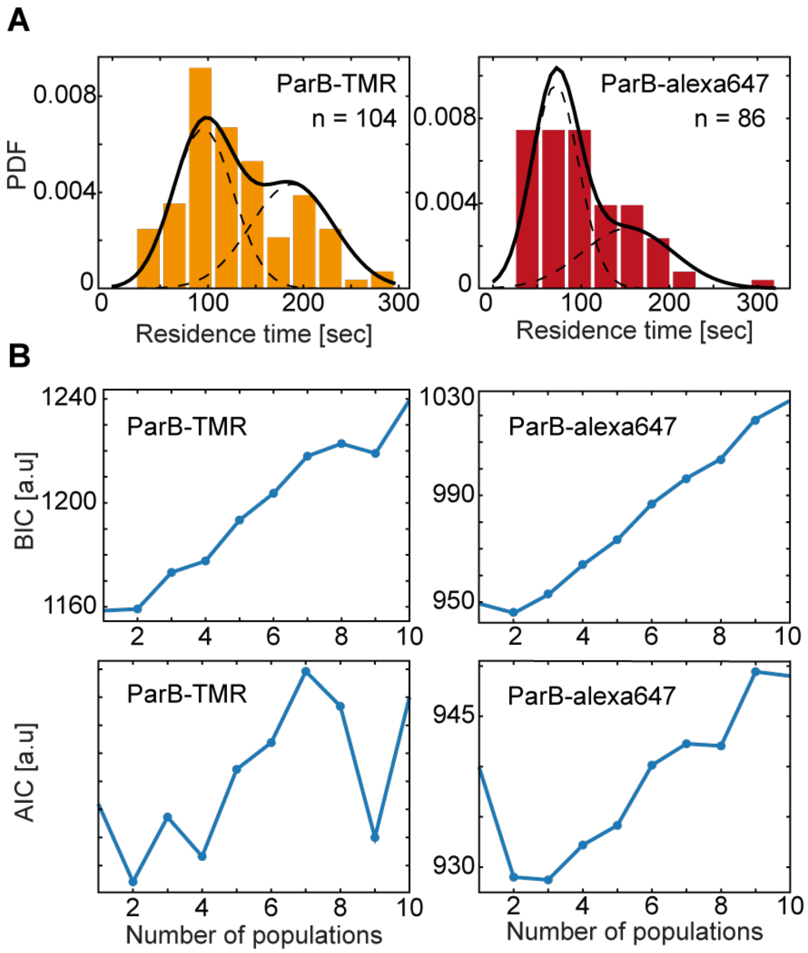


**Figure S3. ParB residence time with different fluorophores. A)** Histogram and normal-distribution fit of the residence times $t_R$ for $ParB^{TMR}$ (left) and $ParB^{alexa647}$ (right). **B)** Bayesian information criterion (BIC, top) and Akaike information criterion (AIC, bottom) for corresponding fits with multiple number of populations. The final fits were chosen at 2 populations, in line with previously reported data (*1*, *5*).

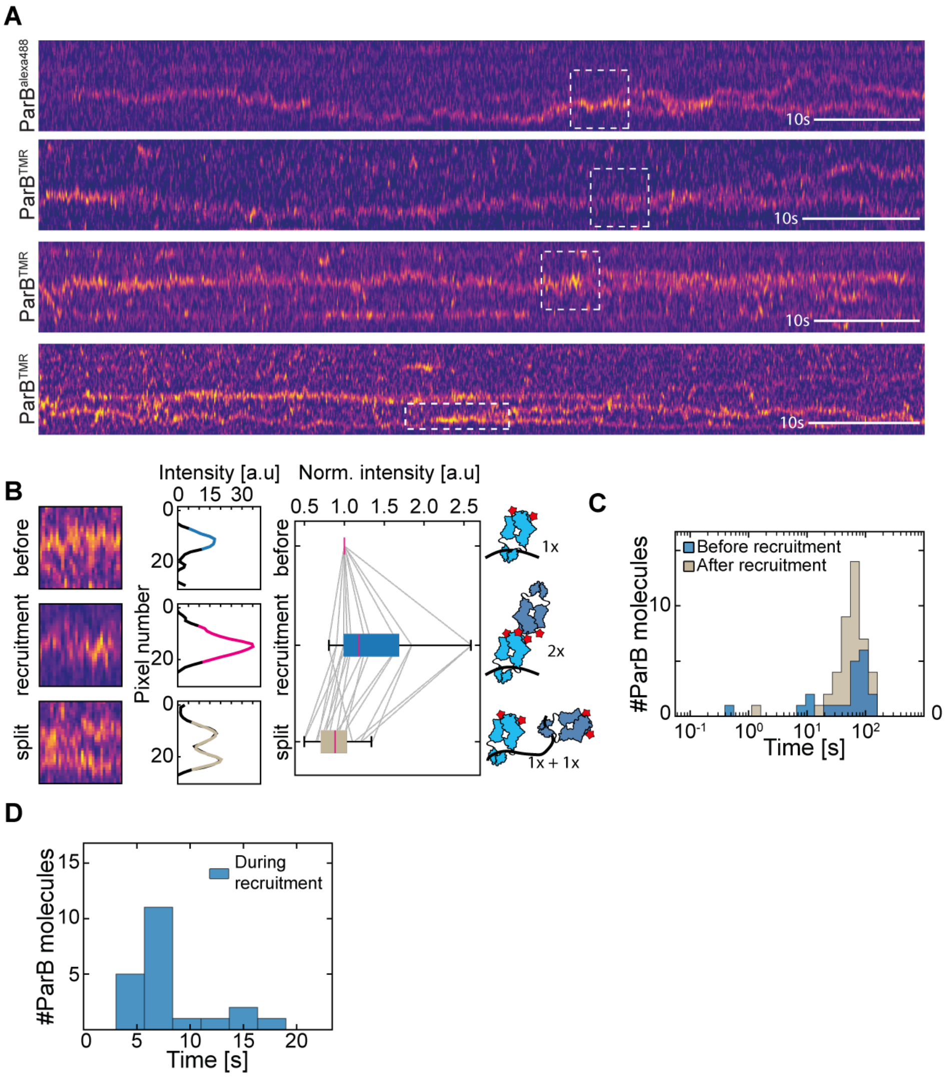


**Figure S4. ParB-ParB recruitment is marked by short conjunct diffusion followed by a split and two individually diffusing ParB dimers. A)** Kymographs for ParB-ParB recruitment using ParB proteins with different tags. Top-to-bottom: ParB-alexa488, 3xParB-TMR. Scale bar = 10s. Dashed boxes represent ParB-ParB recruitment event. **B)** Left: exemplary parts of the kymographs "before recruitment", "during recruitment" and "split after recruitment" from traces such in panel A). Window size is 25px*25 frames. Middle: Mean intensity projection from the windows on the left. Right: Relative intensity "during recruitment" and "split after recruitment", as normalized to the "before recruitment" of the same kymograph. n=25 before and n=50 after splitting. Cartoon representations on the right represent the ParB dimerization that explains the increased signals during recruitment. **C)** Residence time of single ParB dimers before (blue) and after (cream) recruitment events. **D)** Time duration of the conjunct diffusion period during the recruitment event.

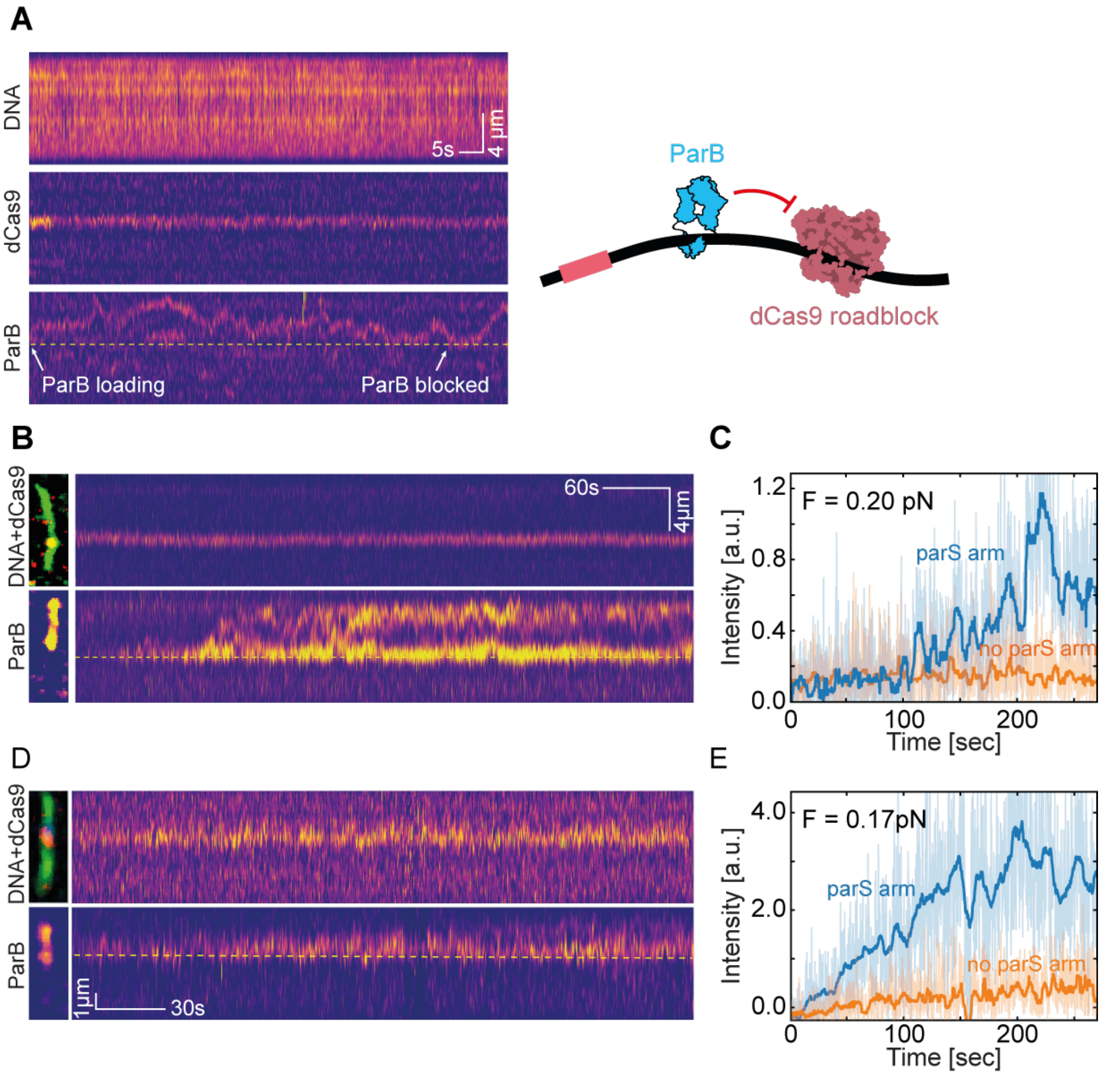


**Figure S5. DNA-bound dCas9 roadblocks efficiently block diffusion of ParB dimers. A)** Kymographs for DNA$_{parS}$ stained with SytoxGreen (top) and dCas$^{alexa549}$ (middle) and ParB$^{TMR}$ (bottom) at low concentration [0.1nM]. White arrows indicate the loading position (left) and the moment of dCas9 blocking ParB diffusion (right). Cartoon representation shown on the right. Scale bars: spatial 4µm, temporal 5s. **B)** Kymographs for dCas$^{alexa549}$ (top) and ParB$^{alexa647}$ (bottom) at 10x higher ParB concentration [1nM]. Left images are snapshots of dCas9$^{alexa549}$ & $^{SytoxGreen}$DNA$_{parS}$ overlay, and ParB$^{alexa647}$ signal at the end of the trace. Yellow dashed line indicates the position of the dCas9$^{alexa549}$. Scale bars: spatial 4µm, temporal 60s. **C)** Quantification of the kymograph data of panel B, that displays the ParB signal in the top (blue, raw data – light blue) and bottom (orange, raw data – light orange) part of the DNA, i.e., above and below the Cas9 (dashed yellow line), respectively, at F = 0.20 pN. **D)** Same as panel B. **E)** Same as panel C.

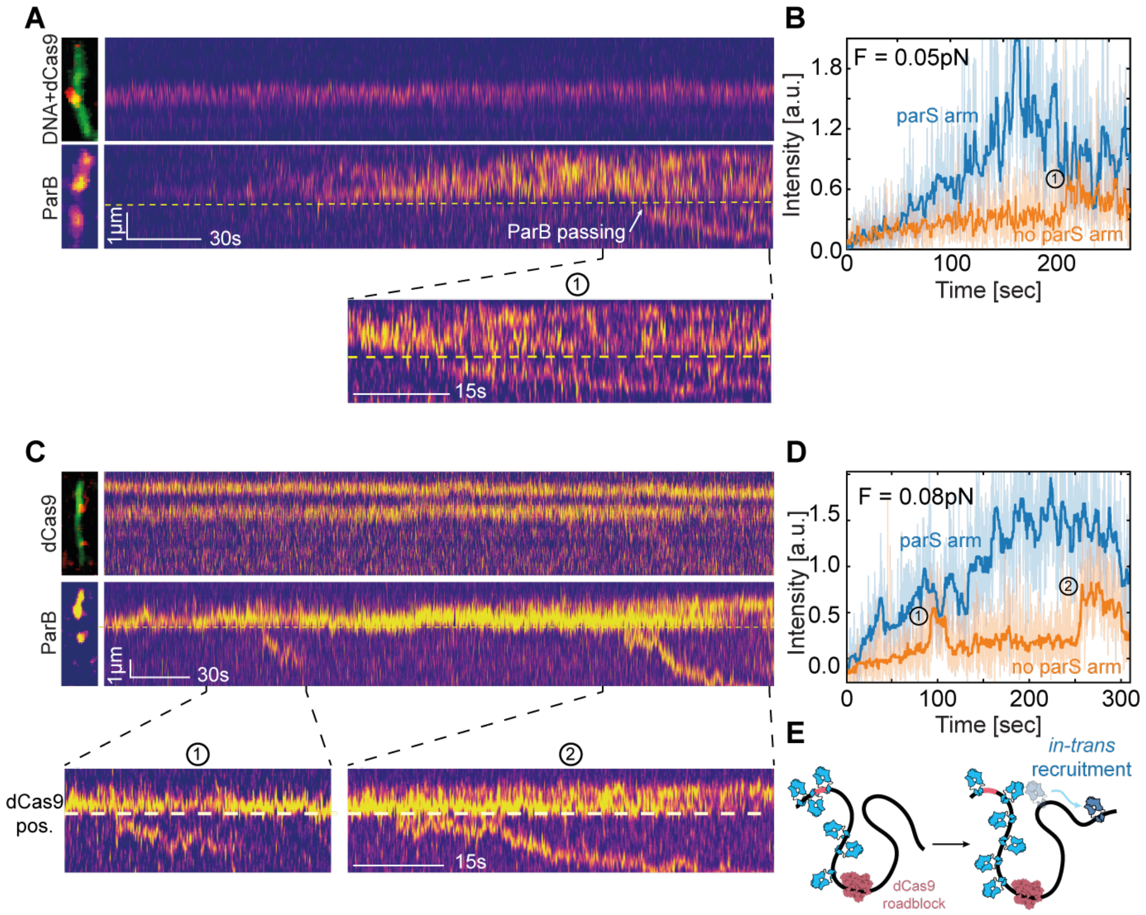


**Figure S6. ParB-ParB in-trans recruitment allows for bypassing DNA-roadblocks at low forces. A)** Kymograph for dCas9$^{alexa549}$ (top), and ParB$^{alexa647}$ (bottom) for DNA$_{parS}$ (same as Fig.S5B) at F= 0.05 pN. Left images are snapshots of dCas9$^{alexa549}$ & $^{SytoxGreen}$DNA$_{parS}$ overlay and ParB$^{alexa647}$ signal at time of recruitment (2). Yellow dashed line indicates the position of the dCas9$^{alexa549}$. Scale bar 30s. Zoomed region (1) represents event where ParB is recruited over the dCas9-roadblock, and continues diffusion over it. **B)** Quantification of the kymograph data of panel A, that displays the ParB signal in the top (blue, raw data – light blue) and bottom (orange raw data – light orange) part of the DNA, i.e., above and below the Cas9 (dashed yellow line), respectively. (1) represents a crossing event where the intensity increases on the 'no *parS* side'. **C)** Same as in panel A. Zoomed regions (1) and (2) represent two events where ParB is recruited over the dCas9-roadblock, and continues diffusion over it. Scale bar = 30s **D)** Same as panel B. (1) and (2) represent crossing events that are marked by an intensity increase on the 'no *parS* side'. **E)** Cartoon representation of the hypothesized ParB-ParB *in-trans* recruitment event.

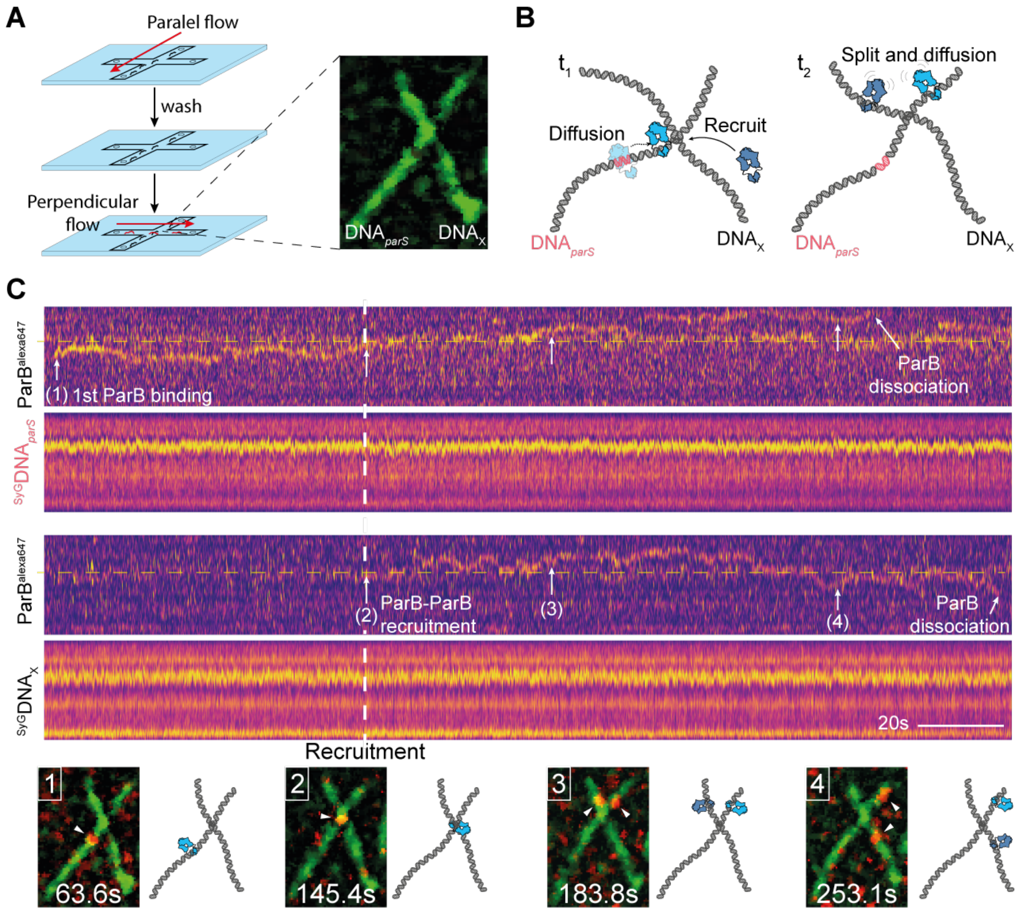


**Figure S7. A new ParB dimer can be recruited by a ParB dimer on a different DNA molecule. A)** Schematic representation of the experimental setup flow cell and simplified experimental procedure (see Methods). Parallel flow is applied to bind a DNA$_{parS}$ molecule to the surface. After washing, perpendicular flow is applied to bind DNA$_X$ molecules at a large angle compared to the initially bound DNA$_{parS}$. Right: single frame snapshot of the final arrangement of DNA$_{parS}$-DNA$_X$. **B)** Cartoon representation of the experiment. *parS*-sequence is indicated in pink **C)** Kymographs for ParB$^{alexa647}$ (top) and DNA stained with SytoxGreen (bottom). Top two kymographs represent the DNA$_{parS}$ signal taken from intensity profiles along the DNA$_{parS}$ molecule. Bottom two kymographs represent DNA$_X$ signal taken from intensity profiles along the DNA$_X$ molecule. Dashed yellow line indicates the approximate junction point. Dashed white vertical line indicates the time when the event occurred where ParB on the DNA$_{parS}$ molecule met the junction point and a new ParB dimer was loaded onto the DNA$_X$ molecule. Bottom row shows single frame snapshots (with corresponding cartoon representations on the side) of the DNA$_{parS}$-DNA$_X$ molecules at timepoints indicated by (1)-(4).

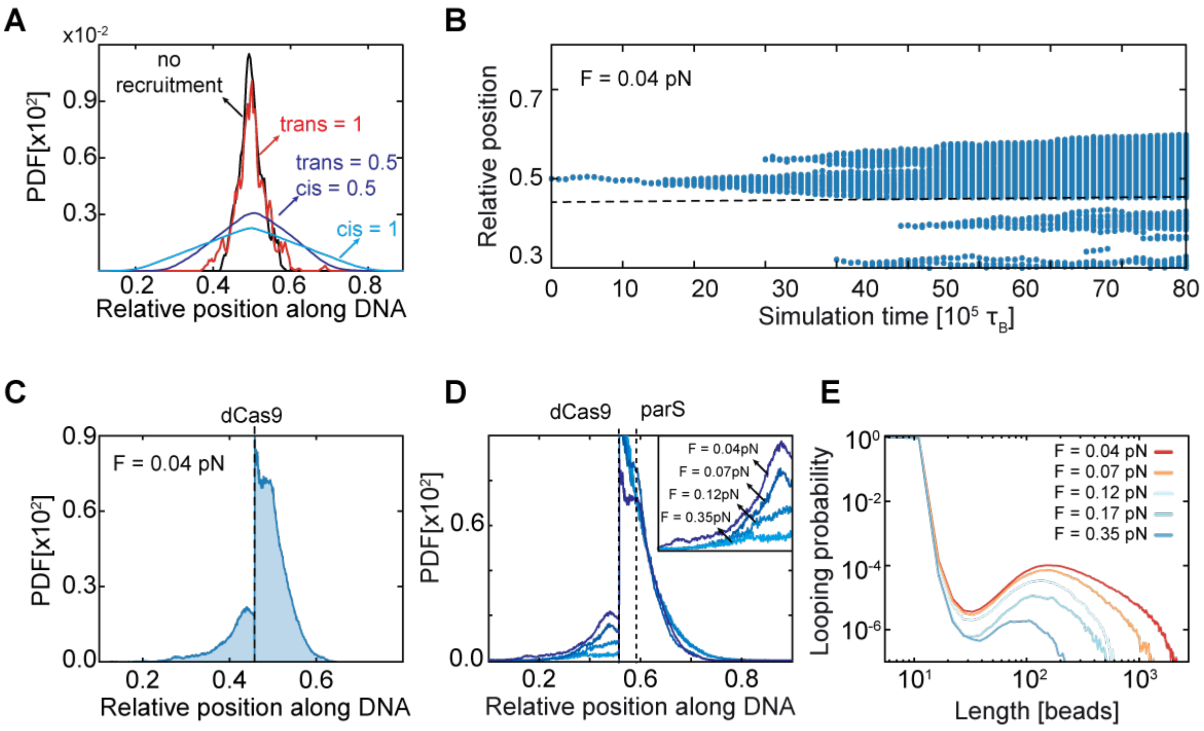


**Figure S8.** ***In-trans*** **recruitment leads to roadblock-passing via DNA looping. A)** ParB spreading with different in-built recruitment ratios (in all cases: *cis*+*trans* = 1, *see Methods*) copmpared to "no recruitment" scenario, equivalent to clamping and spreading model. **B)** Histogram of ParB positions during simulation experiments (F = 0.04 pN). Dashed line indicates the position of the roadblock particle. **C)** Histogram of cumulated ParB probability density from MD simulations averaged over n=64 simulations represented in panel B. Dashed line indicates the position of the roadblock particle. **D)** Same as C, with overlayed histograms at different forces (tether lengths). **E)** Polymer looping probability as the function of tether length (force to length conversion based on WLC model (*6*)**.**

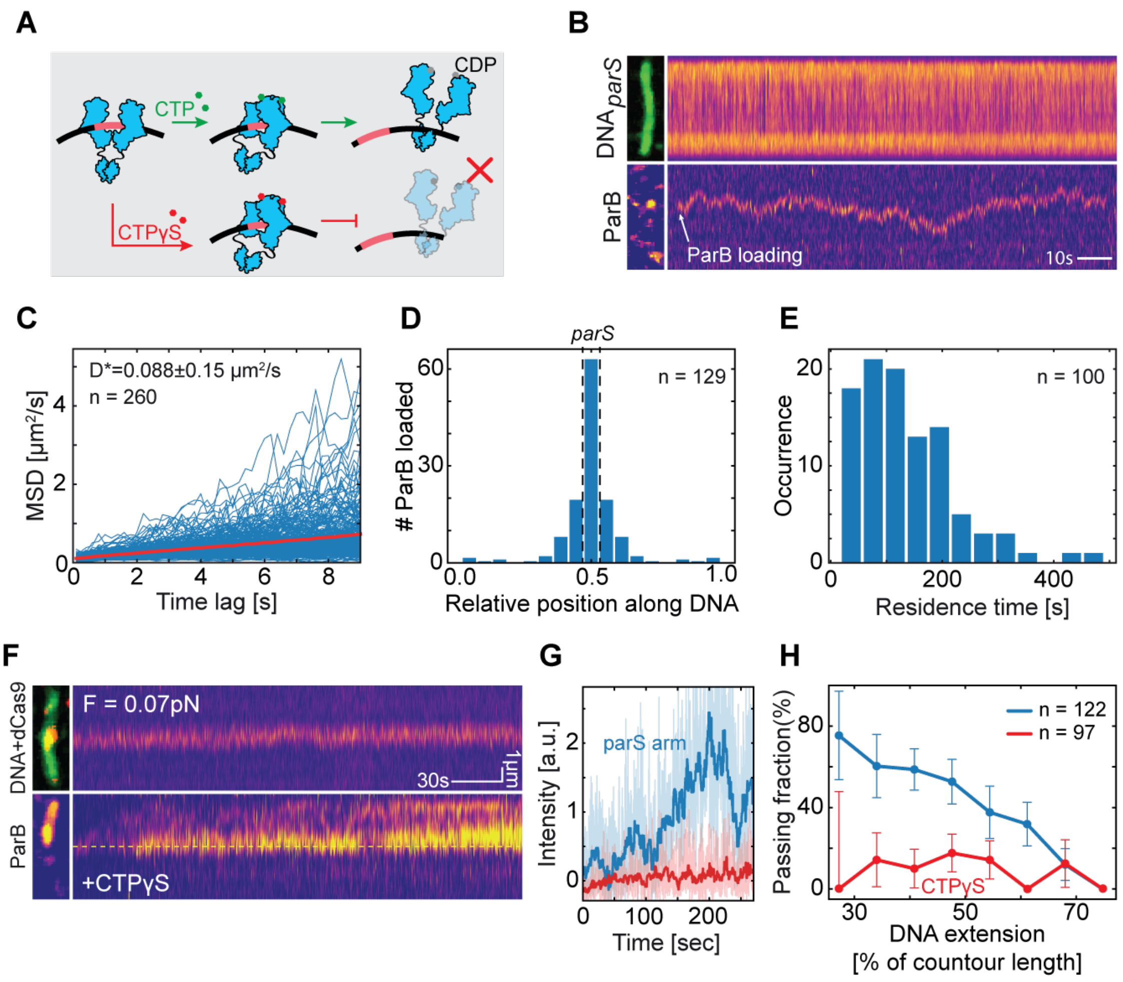


**Figure S9. CTP is not required for spreading but is required for ParB-ParB recruitment. A)** Cartoon representation of ParB in the presence of CTP and CTPγS. While CTP allows for hydrolysis and clamp opening, CTPγS largely prevents that and retains ParB in a closed conformation (*7*, *8*). **B)** Kymographs for $DNA_{parS}$ stained with SytoxGreen (top) and $ParB^{alexa647}$ (bottom). Single frame snapshots of the DNA and ParB at the moment of binding, are provided on the left. White arrow indicates ParB loading. Scale bar = 10s. **C)** Mean square displacement of the diffusing ParB molecules loaded at *parS* site in the presence of CTPγS. Apparent diffusion coefficient is D = 0.088±0.015 μm$^2$/s, n= 260. **D)** Mirrored histogram representing loading position of ParB molecules in the presence of CTPγS relative to the $DNA_{parS}$ ends. *parS* site position is represented by dashed lines. n=129. **E)** Residence times of diffusing ParB molecules after binding the $DNA_{parS}$ in presence of CTPγS (n=100). **F)** Kymograph for $dCas9^{alexa549}$ (top), and $ParB^{alexa647}$ (bottom) for $DNA_{parS}$ at F= 0.07 pN in the presence of CTPγS. Snapshots of $dCas9^{alexa549}$ and $^{SytoxGreen}DNA_{parS}$ overlay and $ParB^{alexa647}$ signal are provided on the left. Yellow dashed line indicates the position of the $dCas9^{alexa549}$. **G)** Quantification of the kymograph data of panel F, that displays the ParB signal in the top (blue) and bottom (red) part of the DNA, i.e. above and below the Cas9 (dashed yellow line), respectively. F = 0.07 pN. **H)** Fraction of DNA molecules that exhibit ParB dimers passing the dCas9-roadblock. Blue and red data are for CTP (blue, n=122) and slowly-hydrolysable CTPγS (red, n=97), respectively(*9*). Error bars represent binomial proportion confidence intervals.
